# Deep Representation Learning Determines Drug Mechanism of Action from Cell Painting Images

**DOI:** 10.1101/2022.11.15.516561

**Authors:** Daniel R. Wong, David J. Logan, Santosh Hariharan, Robert Stanton, Andrew Kiruluta

## Abstract

Fluorescent-based microscopy screens carry a broad range of phenotypic information about how compounds affect cellular biology. From changes in cellular morphology observed in these screens, one key area of medicinal interest is determining a compound’s mechanism of action. However, much of this phenotypic information is subtle and difficult to quantify. Hence, creating quantitative embeddings that can measure cellular response to compound perturbation has been a key area of research. Here we present a deep learning enabled encoder called MOAProfiler that captures phenotypic features for determining mechanism of action from Cell Painting images. We compared our method with both a traditional computer vision means of feature encoding via CellProfiler and a deep learning encoder called DeepProfiler. The results, on two independent and biologically different datasets, indicated that MOAProfiler encoded MOA-specific features that allowed for more accurate clustering and classification of compounds over hundreds of different MOAs.

## Introduction

High-content screening (HCS) produces diverse phenotypic information that is of great interest to the drug discovery process^1,2^. One such procedure known as Cell Painting^3^ allows for broad profiling of cellular phenotypes in response to different compounds. Much effort has gone into quantitatively characterizing these phenotypes^4–6^, with the goal of creating representations for describing how different compounds affect biology. Quantitatively profiling cellular response to compound perturbation has many use cases, such as target identification, concentration optimization, and mechanism of action (MOA) determination (e.g. heat shock protein inhibitor, CDK inhibitor, histamine receptor antagonist)^7^.

Traditional computer vision methods such as CellProfiler (CP)^8^, which is the gold standard in cellular profiling, rely on extracting preset human-selected features from image data. Methods like this have proven useful for encoding subtle changes in cellular phenotype to derive biological insight, with a variety of applications such as object detection^9^, cell viability assessment^10^, and MOA determination^7^. Although traditional computer vision techniques have proved useful, they often require much fine-tuning and require human intelligence and intuition for deciding which phenotypic features and their parameters are important to measure. In contrast, deep learning has emerged as a tool for learning and encoding meaningful representations^4^ (i.e. embeddings) without requiring humans to know beforehand what features may be useful for the task of interest. Indeed, deep representation learning^11,12^ has facilitated improved understanding in both biology^12–14^ and medicine^15–17^.

One goal of phenotypic HCS is to determine the MOAs of compounds^18^. Ascertaining MOA is a challenging but worthwhile task that can provide insight into drug efficacy, side effects, dosing, and possible success in clinical trials^19,20^. Some success has been seen outside of phenotypic screening endeavors, such as through analyzing a compound’s structure and effect on genetic profiles^21–23^. For phenotypic screening, deep learning has performed better than traditional techniques for MOA determination^24–29^. However, these studies only served as a proof-of-concept, encompassing a small set of about a dozen MOAs. Here, we present a MOA determination method spanning hundreds of MOAs that showed efficacy on two independent datasets: 1) the Joint Undertaking in Morphological Profiling (JUMP1) pilot dataset^30^ encompassing 176 MOAs 2) the Library of Integrated Network-Based Cellular Signatures (LINCS) dataset^31,32^ encompassing 424 MOAs. We compared our method called MOAProfiler (MP) with CP as well as a deep learning based method called DeepProfiler (DP)^33,34^. We present MP as an open-source and readily available tool for deep phenotypic profiling of Cell Painting images.

## Results

### JUMP1 Performance

To assess whether we could determine a broad range of MOAs from phenotypic information alone, we developed a deep learning method across two datasets differing in cell type, concentration, and time-points (Methods). For the first, we turned to the JUMP1 dataset which had MOA metadata information provided by the Clue Connectivity Map^35^. To simplify the compound space and limit the intricacies associated with polypharmacology^36^, we took a subset of the compounds that had at most one known MOA. The resulting dataset spanned 266 compounds and 176 MOAs (Figure 1A). Most compounds were plated at 23 replicate wells (Figure 2A, left) and most MOAs were represented by one compound (Figure 2B, left). There were either 9 or 16 image fields per well (Figure 2C, left). Cell Painting images consisted of five channels capturing different areas of cellular morphology (Figure 1B).

**Figure 1:**
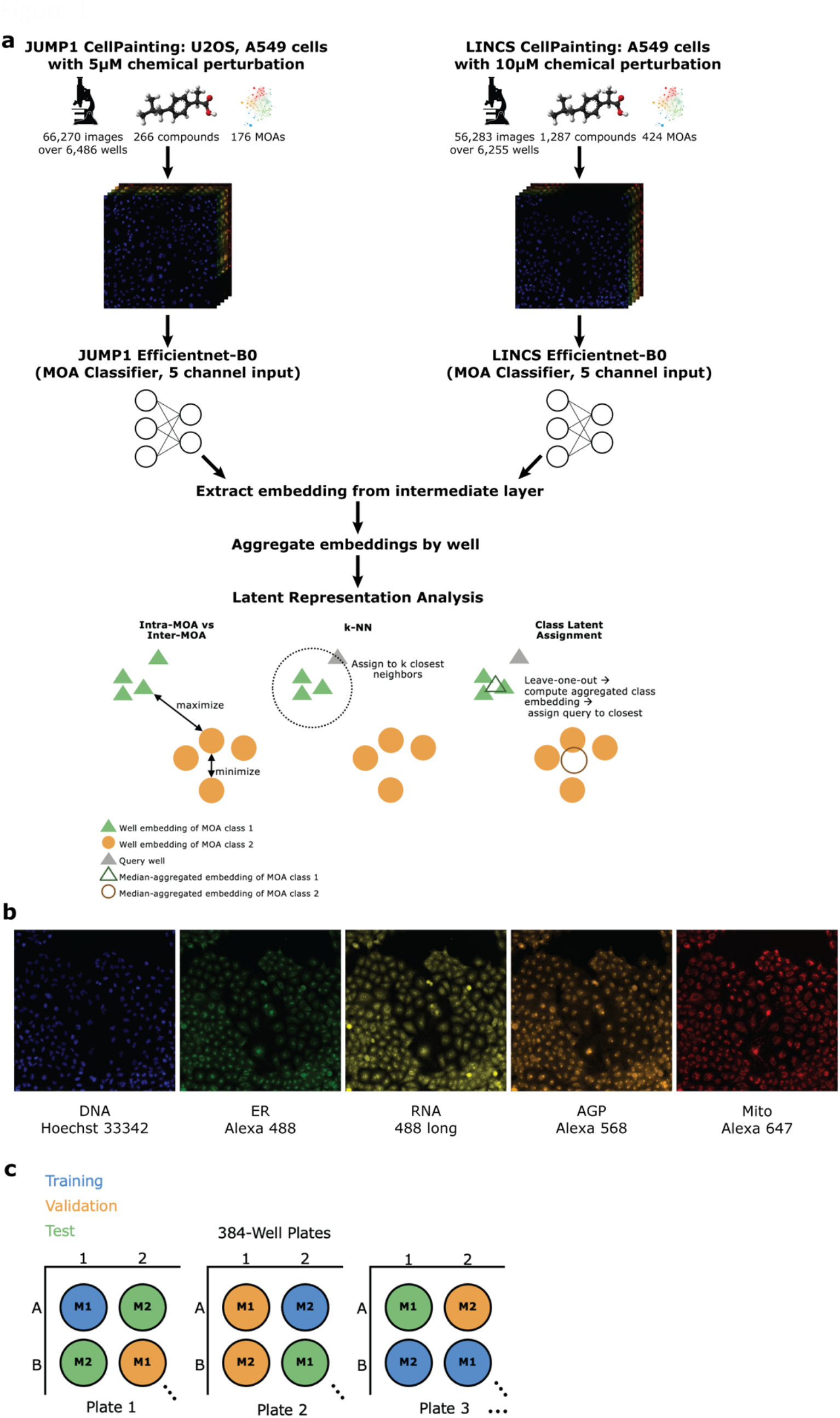
Study overview. a) Study overview. We applied the same method to two datasets independently: the JUMP1 Pilot dataset as well as the LINCS dataset. Training and analysis were performed separately for each study. Below: pictorial visualizations of the various metrics assessing embedding viability and clustering. b) Example Cell Painting images from the JUMP1 dataset. Color added for visualization. DNA = deoxyribonucleic acid, ER = endoplasmic reticulum, RNA = ribonucleic acid, AGP = actin cytoskeleton, Golgi, plasma membrane, Mito = mitochondria. c) Schematic of training, validation, test split. M1 indicates MOA class one and M2 indicates MOA class two. Each circle is a well. The schematic is for the sake of illustration and is not the actual location split used in the study.

**Figure 2:**
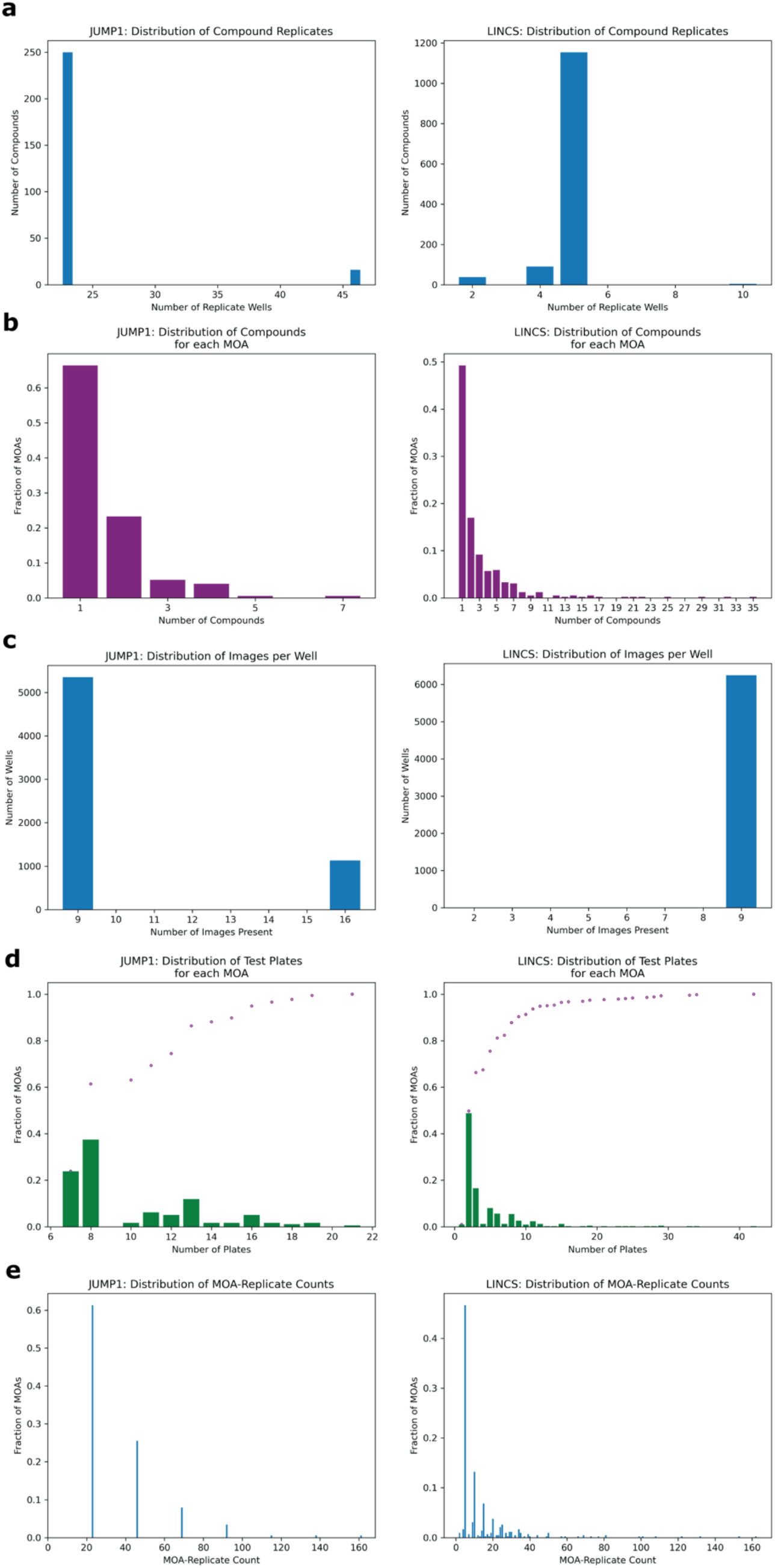
Distributions of study metadata. All panels exclude the negative control DMSO condition. JUMP1 (left), LINCS (right). a) Distribution of replicate well count per compound. E.g. left panel: Most compounds were each plated in 23 wells. b) Distribution of compounds for each MOA. E.g. left panel: 66% of MOAs were represented by one compound each. c) Distribution of images per well. Most wells had 9 non-overlapping image fields. d) Distribution of number of plates in the test set for each MOA. Example from left figure leftmost bar: 24% of MOAs were each represented on 7 plates in the held-out test set. Purple dots show the cumulative frequencies at each plate count. e) Distribution of MOA-replicate counts. “MOA-replicates” in this case is defined as any wells with any compounds with the same MOA. E.g. left panel: 62% of MOAs were each represented by 23 different wells in the entire dataset.

To develop a model capable of creating MOA-specific embeddings from cellular phenotype, we trained an EfficientNet model^37^ to classify an image field’s MOA, with the motivation of directly learning which features are important for segregation. Only images were provided as input to the model, with no information on compound, concentration, or any other form of metadata (Methods). To minimize the model learning confounding experimental features like well-location specific anomalies, we split the dataset such that each well was randomly assigned to only one of three sets: training, validation, or test (Figure 1C). We divided the dataset such that 60% of the wells were assigned to training, 10% to validation, and 30% to test (Methods). We also ensured each MOA’s test wells spanned multiple plates (at least seven, Figure 2D, left) to assess the model’s performance across potential variation of plate-specific conditions. Since MOA-replicate count was imbalanced (Figure 2E, left), we measured the model’s precision recall characteristics. On the held-out test set, the model achieved an area under the precision recall curve (AUPRC) of 0.46 (random AUPRC = 0.006) for image field classification over 176 MOA classes (Figure 3A). This equated to an accuracy of 0.52 (random accuracy = 0.006). To assess the influence of well-location effects which are common in microscopy^38,39^, we compared the model’s performance on edge wells versus non-edge wells. We found minor differences in classification accuracy (0.54 vs .50) suggesting that the model was not leveraging much confounding edge-specific features for its learning (Figure 4).

**Figure 3:**
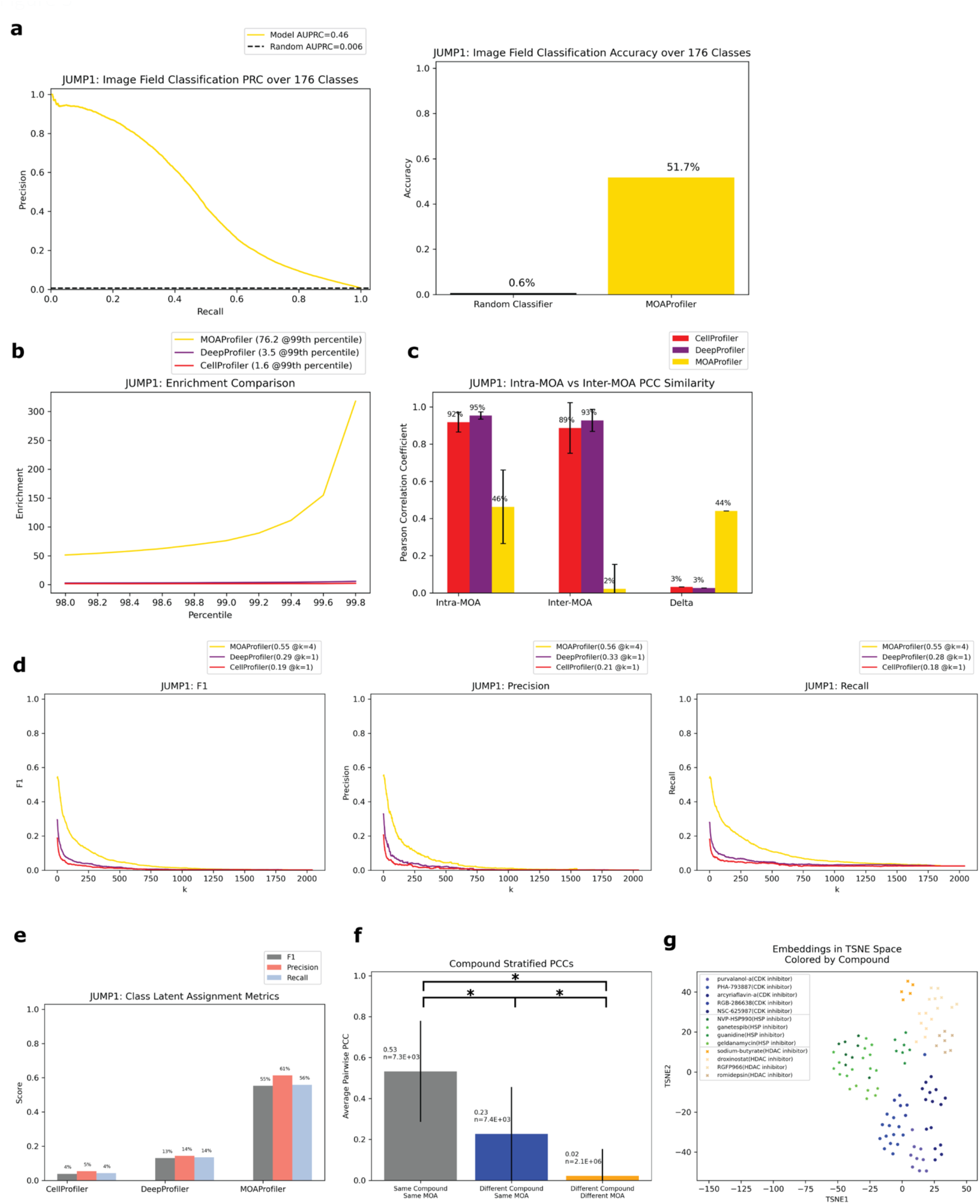
Performance of model trained and evaluated on the JUMP1 Dataset. All evaluation was over the test set. a) Left: PRC of the classifier, random baseline = positive prevalence of binarized labels. Right: classification accuracy, random baseline = 1 / 176. Classification is over raw image fields (not embeddings as in panels d and e). b) Enrichment scores for the three methods from the 98^th^ percentile to the 99.8^th^ percentile with step sizes of 0.2. Enrichment at the 99^th^ percentile (rounded to the nearest tenth) is shown in the figure legend. c) Average of pairwise PCCs for two groups: Intra-MOA (embeddings with the same MOA class), Inter-MOA (different MOA classes). Delta = Intra-MOA average - Inter-MOA average. Error bars span one standard deviation in each direction. d) k-NN metrics for embedding classification calculated for all values of k. F1 (left), precision (middle), and recall (right). The highest score for each method and corresponding k are shown in the legend (rounded to the nearest hundredth). e) F1, precision, and recall values for the class latent assignment metric. Scores rounded to the nearest hundredth. f) Average of pairwise PCCs for three groups of embeddings. For each group, we calculated PCCs for each possible pair of wells. Significance (*) indicates p<<0.0001 for a two-sided z test. Error bars span one standard deviation in each direction. g) TSNE visualization of well embeddings of three example MOAs (chosen because they were each represented by four or more compounds). Circles = CDK inhibitor, stars = HSP inhibitor, x marks = HDAC inhibitor. Different compounds with the same MOA were given similar but different colors.

**Figure 4:**
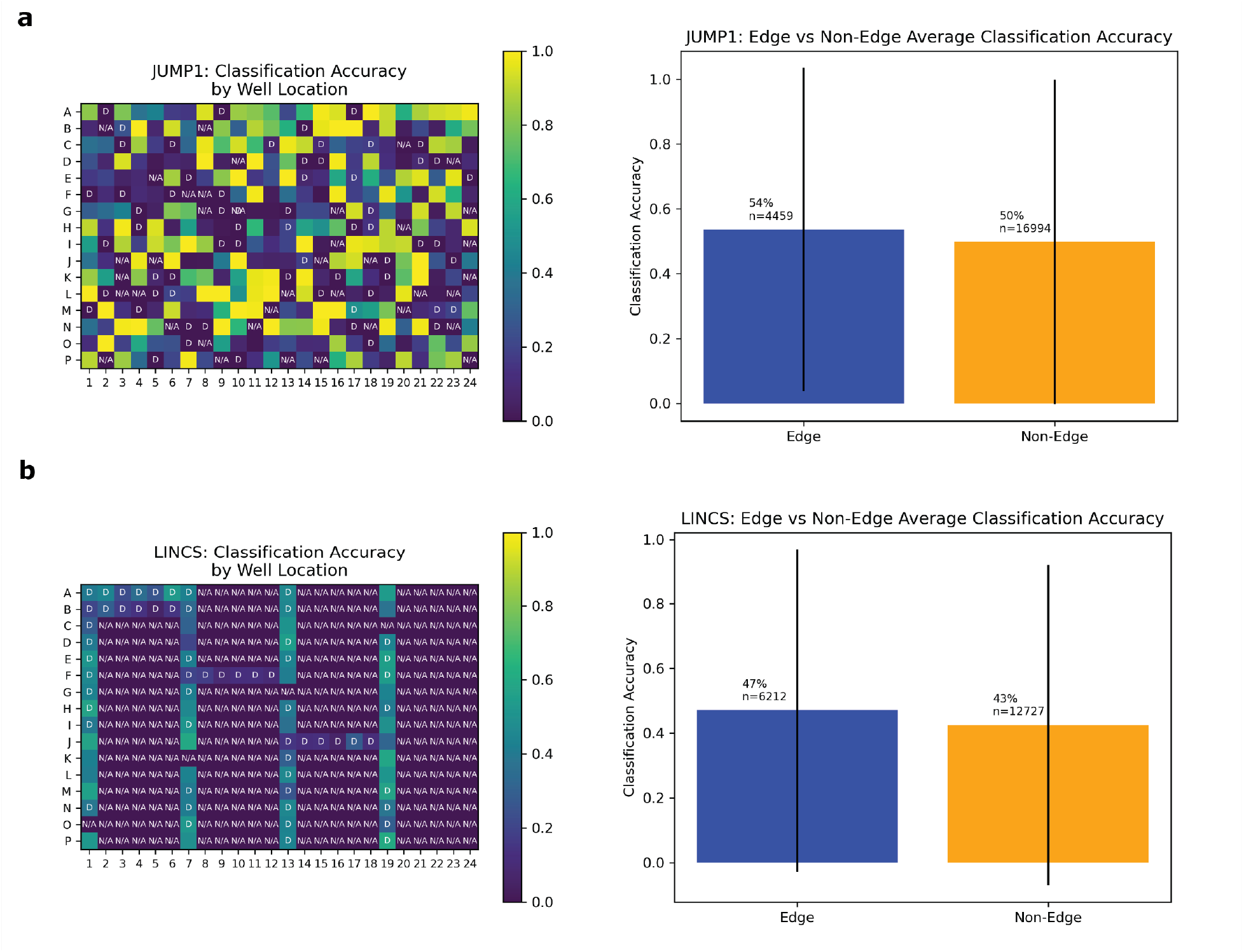
There was a small difference in image classification accuracy for edge wells versus non-edge wells. All evaluation was over the test set. An edge well is a well with rows “A” or “P”, or columns 1 or 24. a) Well location effects for JUMP1 Dataset. Left: 384-well plate heatmap (over 23 plates) of model classification accuracies for image fields. “D” indicates negative control DMSO wells (no perturbation). “N/A” indicates wells excluded in our study due to the compound having more than one known MOA. Right: Average classification accuracy for images (not embeddings) residing in edge and non-edge wells. Accuracies were calculated over all image fields with sample size “n”. b) Well location effects for LINCS Dataset. Same analysis as in (a) except the LINCS Dataset was over 136 384-well plates. “N/A” indicates excluded wells where compounds had more than one known MOA or were not at 10μM.

With the trained classification model, we measured how well the model created embeddings that were meaningful for MOA classification. This differed from simply using the model as a classifier, which would have limited use outside of the fixed MOA set on which it was trained. Hence, we extracted image embeddings from an intermediate layer in the network^12^, aggregated them by well by taking the median, and assessed how valuable these well-level embeddings were for the task of MOA classification compared to CP and DP (Methods).

Ideally, embeddings with the same MOA (intra-MOA) should be more similar than embeddings with different MOAs (inter-MOA). Hence, we measured how likely it was to observe intra-MOA embeddings in strongly correlated versus weakly correlated embedding pairs. Through a Fisher’s exact test, we found that intra-MOA embedding pairs were more likely to be found in strongly correlated versus weakly correlated embedding pairs, with greater enrichment of MP-derived embeddings vs CP-derived and DP-derived embeddings (enrichment at the 99^th^ percentile = 1.6 CP, 3.5 DP, 76.2 MP, Figure 3B). Similarly, we asked which of the methods could better generate embeddings that captured phenotypic differences between the MOAs. There was greater difference between intra-MOA embeddings and inter-MOA embeddings when constructed by MP instead of CP or DP (delta = 0.44 for MP, Figure 3C). For both CP and DP, the difference was smaller (delta = 0.03 for CP, 0.03 for DP). This indicates that MP encoded different MOAs in different and distinguishable phenotypic spaces, which may be advantageous if a new compound were to be queried for its MOA.

To simulate this situation of predicting the MOA of a query compound using its phenotypic embedding, we performed two analyses: k-nearest-neighbors (k-NN) and an analysis we constructed called the “class latent assignment” (Figure 1A for a pictorial visualization, Methods). We used each well in the test set as a held-out query. For k-NN, we predicted the query well’s MOA as the majority MOA of its k-nearest neighbors (Methods). For all values of k, we calculated F1, precision, and recall values. We found that for all metrics, MP outperformed CP (percent improvement: 191% F1, 170% precision, and 202% recall). MP also outperformed DP (percent improvement: 85% F1, 69% precision, and 96% recall, Figure 3D).

The “class latent assignment” method was a parallel way to classify a query compound’s MOA by instead using similarity to aggregated MOA-class embeddings rather than to well-level embeddings for predicting a query well’s class (Methods). This metric has the advantage of being less sensitive to single-well outliers, and reduces the impact of the immediate closest neighbors so that embeddings can be queried against more representative class-wide embeddings. We computed an aggregated MOA-level embedding (MLE) for each MOA by taking the median of all the same-MOA well embeddings with the query well’s embedding left out. We then predicted the query well’s MOA as the MOA of the most similar MLE. We calculated F1, precision, and recall scores for the resulting predictions. By this metric, MP was more performant than CP (percent improvement: 1333% F1, 1050% precision, and 1228% recall). MP also exceeded DP’s metrics (percent improvement: 325% F1, 327% precision, and 314% recall, Figure 3E).

Since some compounds have the same MOA, we wanted to know whether the model was learning MOA-specific phenotypes consistent through different compounds with the same MOA, as opposed to simply learning each compound’s phenotypic effect. Hence, we calculated the average Pearson Correlation Coefficient (PCC) of three groups: well pairs with 1) the same compound (and hence same MOA) 2) different compounds with the same MOA and 3) different compounds with different MOAs. We found that of the embedding pairs with different compounds, the pairs with the same MOA class were more similar than those with different MOA classes (average PCC = 0.23 vs 0.02, Figure 3F). The three populations’ PCC averages were all significantly different (p<<0.0001 for all two-sided z-tests). This suggests that indeed the phenotypic embeddings that the model encoded were MOA-specific rather than compound-specific. From a low-dimensional t-distributed stochastic neighbor embedding (TSNE) visualization of embeddings from three example MOAs, we could see that different compounds with the same MOA were clustered together with different MOAs inhabiting different areas in latent space (Figure 3G).

Since most MOAs were only represented by one compound each (Figure 2B, left), we also assessed the performance metrics with a subset of the test dataset that only included MOAs that were each represented by multiple compounds. In this analysis, we wanted to assess whether performance was over-inflated by single-compound MOAs and whether the model had simply learned to group compound replicates, which could be the case if the dataset was heavily over-represented by single-compound MOAs. However, the model created embeddings that were clustered by MOA despite each MOA being represented by multiple compounds (Supplemental Figure 1).

To simulate the real-world use case of identifying MOAs of unknown held-out compounds, we performed an analysis where we split the dataset by compound instead of by wells (Methods). In this scheme, we randomly selected and held out one compound for each of the MOA classes that were represented by at least two distinct compounds. We chose a threshold of two compounds so that each held-out compound would have at least one other same-MOA compound in the training set to facilitate learning. This resulted in 59 compounds that we used as a test set, with the remaining 207 compounds (plus negative control DMSO) used for training and validating a new model (Methods). Despite the model never being exposed to these held-out compounds during training, it was able to correctly predict MOAs for 10.2-13.6% of the compounds in a space of 176 possible MOAs. Compared to a random baseline of 0.6%, this is a 17.9-23.9x improvement. Performance varied depending upon whether we predicted MOA by the neural network’s classification output or by a compound-level embedding’s (CLE) similarity to MLEs of the training compounds (Figure 5).

**Figure 5:**
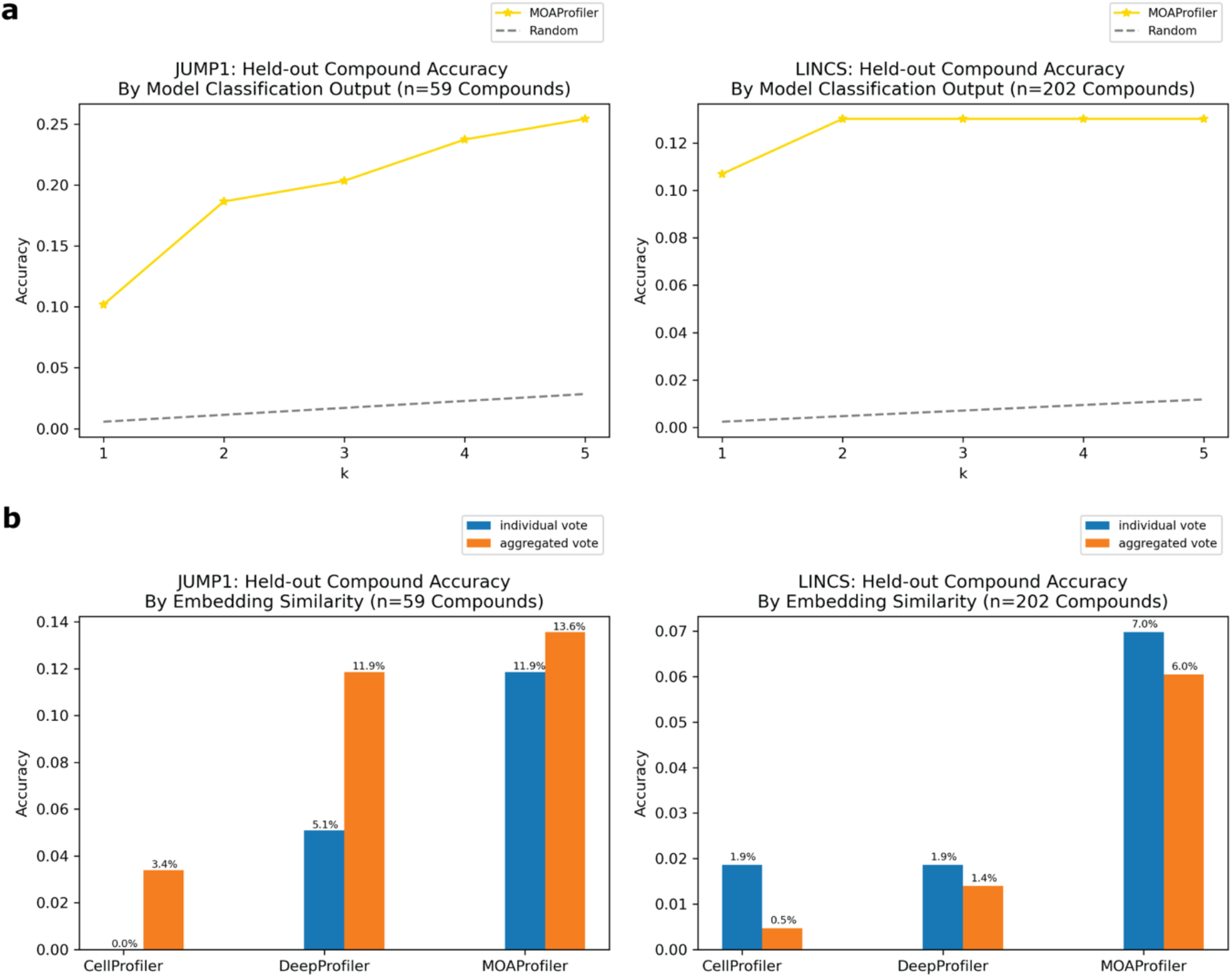
Performance on held-out compounds via two methods. Left: JUMP1, right: LINCS. Y-axis: fraction of held-out compounds that were correctly predicted. a) We used the model’s final image classification output to derive MOA predictions (Methods). “K” is the number of top-MOA predictions that were compared with the true label (i.e. if the true MOA label was in the top-k MOAs, this was counted as a correct prediction). Random baseline is k / number of possible MOAs. b) We used embedding similarities to the training compound embeddings to derive MOA predictions (Methods). Individual vote is the scheme in which the MOA is predicted by taking the majority MOA over each well-level prediction. Aggregated vote is the scheme in which we aggregated the held-out compound’s well-level embeddings into a CLE and predicted its MOA as the MOA of the most similar MLE from the training set. We used an exact match with no flexibility (k=1).

### LINCS Performance

To assess the method on a second dataset, we turned to the LINCS Cell Painting Dataset^32^. Like the JUMP1 dataset, we took a subset of the LINCS data that only had compounds with one known MOA. Furthermore, we took a subset of the data at a fixed concentration of 10μM since this dose produced the strongest phenotype across compounds^39^. The resulting dataset spanned 1,287 compounds and 424 MOAs (Figure 1A). Most compounds were plated in five replicate wells (Figure 2A, right) and most MOAs were represented by one compound (Figure 2B, right). There were 9 images per well (Figure 2C, right).

We trained a separate EfficientNet on this dataset with the same task of classifying MOAs from solely image data. We used the same training-validation-test split scheme by well as we did for JUMP1 along with the same hyperparameter choices (Methods). The model achieved an AUPRC of 0.35 (vs .002 random) and an accuracy of 0.45 (vs .002 random) for image field classification over 424 classes (Figure 6A), which is more than twice as many classes as the JUMP1 dataset’s 177 MOAs. Like the JUMP1 dataset, we did not detect much edge-location confounders affecting classification performance (0.47 edge accuracy vs 0.43 non-edge accuracy, Figure 4B).

**Figure 6:**
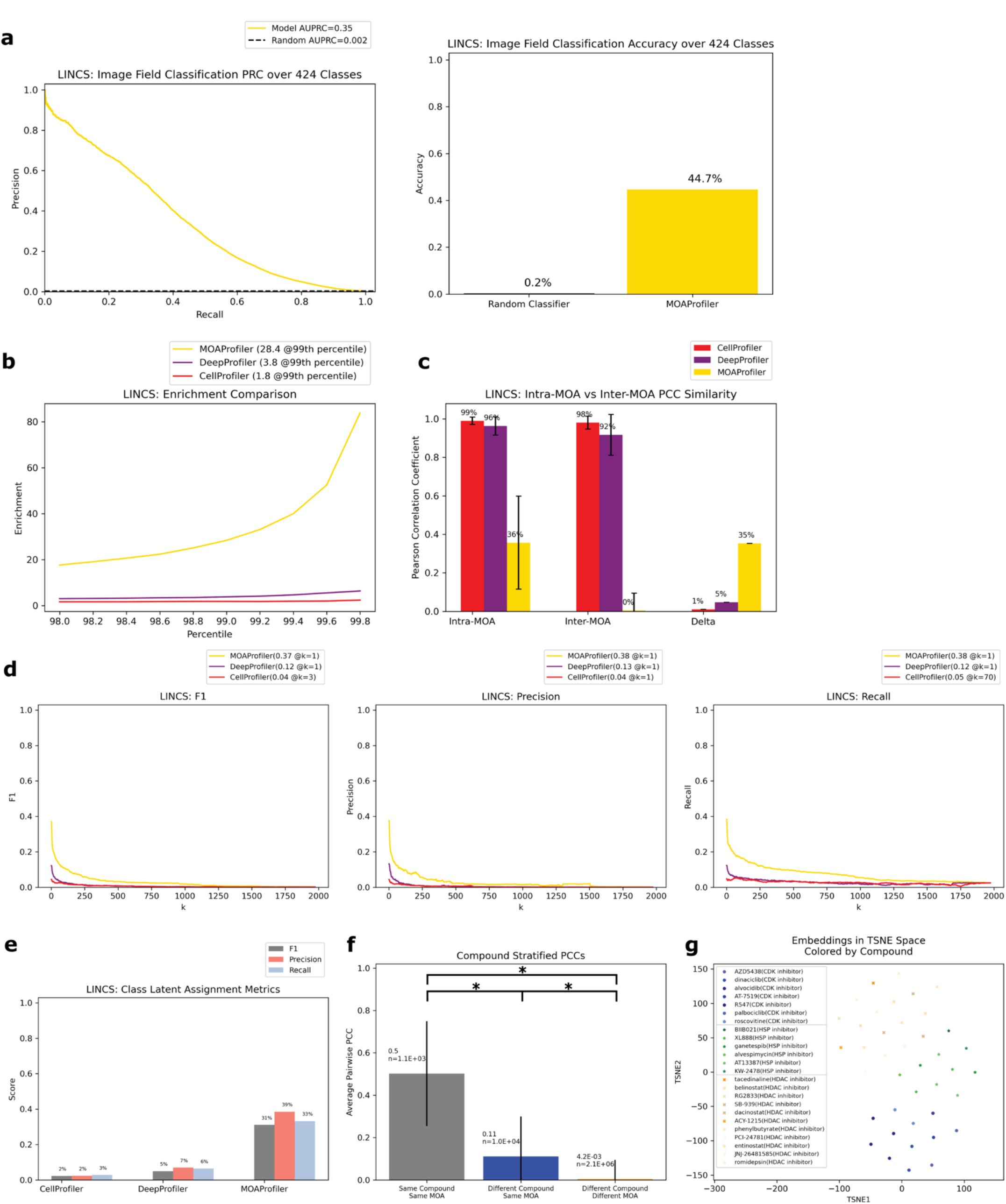
Performance of model trained and evaluated on the LINCS Dataset. Same analysis as Figure 3. All evaluation was over the test set. a) PRC and accuracy of image classification. Random baseline for PRC is 1 / 424. b) Enrichment comparison at different percentiles. c) Intra-MOA vs inter-MOA average pairwise PCC similarity. d) K-NN embedding metrics. e) Class latent assignment metrics. f) Average pairwise PCCs of three different groups stratified by perturbation. TSNE visualization of well embeddings from three example MOAs. Circles = CDK inhibitor, stars = HSP inhibitor, x marks = HDAC inhibitor. Different compounds with the same MOA were given similar colors.

We derived well-level embedding metrics for the held-out test set and determined that MP was advantageous over CP and DP for MOA determination. MP enabled greater enrichment for intra-MOA pairs among strongly correlated embeddings, with 15.5x fold increase in enrichment over CP and 7.4x fold increase over DP (Figure 6B). MP also enabled greater difference between intra-MOA and inter-MOA similarity (delta = 0.01 CP, 0.05 DP, 0.35 MP, Figure 6C). MP also facilitated greater k-NN metrics than CP (percent improvement: 730% F1, 764% precision, and 609% recall). We also observed sizeable performance gains compared with DP (percent improvement: 204% F1, 185% precision, and 211% recall). For the class latent assignment metrics, MP outperformed CP (percent improvement: 1286% F1, 1532% precision, and 1074% recall) and DP (percent improvement: 527% F1, 451% precision, and 420% recall, Figure 6E). Like the JUMP1 Dataset, we also observed significantly greater embedding similarity among different compounds with the same MOA (0.11 average PCC) versus different compounds with different MOAs (near zero average PCC) (Figure 6F). This was a smaller differential than what we observed in the JUMP1 dataset, with greater PCC profile variability in the LINCS dataset between different compounds and lower PCC similarity on average between different compounds having the same MOA. From a TSNE visualization of three example MOAs colored by compound, we can see both MOA separability in latent space and clustering between different compounds with the same MOA (Figure 6G).

Like the JUMP1 study, most MOAs were only represented by one compound each (Figure 2B, right). Hence, we also assessed the performance metrics with a test dataset that only included MOAs that were each represented by multiple compounds. Performance here was similar to the performance in the full test set of MOAs. The model created embeddings that were clustered by MOA despite coming from multiple compounds (Supplemental Figure 2).

Like the JUMP1 study, we also performed an analysis where we split the dataset by compound instead of by wells, resulting in 202 compounds that we used as a held-out test set. Despite the model never being exposed to these compounds during training, it was able to correctly predict MOAs for 6.0-10.7% of the compounds in a space of 424 possible MOAs. Compared to a random baseline of 0.2%, this was a 25.6-45.4x improvement. Performance varied depending upon whether we predicted MOA by the neural network’s classification output or by a compound’s latent similarity to training compound embeddings (Figure 5).

## Discussion

Our findings are consistent with the growing body of literature suggesting Cell Painting can capture broad phenotypic changes encodable by deep learning. Two points merit emphasis: 1) We developed an embedding encoder for accurate MOA identification of compounds 2) The approach outperformed both a traditional (via CP) and a deep learning enabled method (via DP) for this task on two independent datasets. Higher similarity of intra-MOA embeddings (even across different compounds) than inter-MOA embeddings suggests that the model was capturing MOA-specific phenotypic features vs simply learning an individual compound’s phenotypic effect. This specificity towards MOA is important because in drug discovery campaigns many compounds could have the same MOA. Furthermore, a compound’s MOA is often unknown. Hence, when trying to ascertain an unknown compound’s MOA, a reasonable hypothesis is to predict its MOA as the MOA of its most phenotypically similar compounds that are known. Since the embedder can encode similar embeddings for different compounds with the same MOA (Figure 3F and 6F), perhaps it can likewise be useful for MOA discovery across a diverse compound space. Indeed, the held-out compound analysis in Figure 5 suggests that the method can be used to identify the MOAs of new compounds the model has never used for training.

Furthermore, when we restricted the analysis to only MOAs represented by multiple unique compounds (Supplemental Figures 1 and 2), we observed similar performance to the case that included all MOAs regardless of compound representation (Figures 3 and 6 respectively). This reinforces the notion that the model had indeed learned shared MOA phenotypes that persisted across different compounds.

The ranking of PCC similarity for the three pairwise-embedding groups (Figures 3F and 6F) fit with our expectation: Same-compound same-MOA > different-compound same-MOA > different-compound different-MOA. However, the average pairwise PCC for the same-compound same-MOA group was lower than expected (average PCC=0.51 for JUMP1, 0.50 for LINCS), indicating embedding variation even between compound replicates. This could be due to several factors, such as natural experimental variation within compound replicates, or the presence of features that are not important for discriminating MOAs (and hence room for model improvement). The average pairwise PCC score of 0.23 (and 0.11 for LINCS) for the different-compound same-MOA pairings was also similar to other recent findings^39^. This indicated that although the model correctly grouped different compounds that had the same MOA into the same class, same-MOA embeddings had diversity. Perhaps this is because different compounds in the same MOA class may have differing degrees to which they induce phenotypic changes, with some compounds exerting more evident changes while others were perhaps more modest, which would then lead to phenotypic embeddings with higher variance. Still, the fact that the average PCC for the different-compound same-MOA group was 12 times higher (and 26 times higher for LINCS) than the different-compound different-MOA average PCC indicates that the embeddings were capturing MOA-specific phenotypes despite the diversity within same-MOA groupings. It is also worth noting that high intra-MOA PCC (like with CP and DP, Figures 3C and 6C) is not necessarily better for the task of discriminating MOAs, especially if the inter-MOA PCC is also high. Hence, although intra-MOA PCCs are lower for MP, it is still better suited for discriminating MOAs.

Accurate performance on test wells never seen during the model’s training suggests that the model was not leveraging well-specific features for its learning, which can be a confounder in microscopy^34^. Furthermore, we ensured that each MOA’s test set wells were spread across multiple plates (Figure 2D). For the JUMP1 data the model did not seem to encode large batch effects^40^ into the embeddings (with differences in average PCC of embeddings around 0.04 between different plates), but this effect was more prevalent for the LINCS data (Supplemental Figure 3). As a future avenue to explore, perhaps preprocessing images to remediate batch effects in the LINCS dataset prior to learning would enhance performance.

The model encoded MOA-associated phenotypes for two independent Cell Painting datasets, and exceeded performance of two other methods for generating MOA phenotypic embeddings. As an alternative to hand-engineered feature extraction, deep learning did not require biological expertise for deciding which features to extract, yet both DP and MP consistently yielded better results than CP. In comparison to CP, MP was more performant for both class latent assignment and k-NN (Figure 3, Figure 6). Since we demonstrated that features specific to the biological domain of interest can be learned, we advocate for deep representation learning over traditional feature engineering for phenotypic MOA determination.

Although DP is another deep learning based approach to phenotypic profiling that also uses an EfficientNet backbone architecture, we observed larger performance gains with MP. This indicates that architecture alone is not sufficient for better MOA profiling, but rather attention to parameters such as learning objective and image scale is necessary (Supplemental Figure 4A). It is also interesting to see MP’s advantage even when compared to a separate DP model that was trained directly on the LINCS dataset instead of on ImageNet (Supplemental Figure 4). This DP model was trained with the objective of classifying compounds as an auxiliary task for the primary task of MOA classification (i.e. weak supervision). When both models were allowed to train on Cell Painting images, MP still yielded improvement in enrichment (5.1x improvement over DP), k-NN metrics (percent improvement: 136% F1, 131% precision, 142% recall), and the class latent assignment metric (percent improvement: 204% F1, 221% precision, 178% recall). DP trained on LINCS had higher performance than DP trained on ImageNet, indicating that feature extraction is most performant when the model is trained on the same data types. Other than the quantitative advantages in performance, MP has a practical advantage over DP of not requiring single cell locations as input, extraction of which can be cumbersome^41,42^. MP’s standing as compared to DP for biological areas of interest other than MOA have not been determined. Indeed DP (or CP) may be a better choice for other biological domains or other datasets.

It is important to note that the metrics of success were confined mainly to the task of identifying MOAs via representation learning. The reason we emphasize proper embedding generation as opposed to accurate image classification is because a classification model is constrained to only the MOAs it was trained with. If a compound’s MOA is not part of the training set, then the classifier will be of little value. In contrast, an embedder can generate representations for any Cell Painting images regardless of MOA (even MOAs not included in the training set). Tests for other biological areas of interest other than MOA, such as cellular viability, drug toxicity, or protein target identification, have yet to be studied. In such cases, deep learning can provide the advantage of learning features that are directly relevant to the biological area of interest (vs undirected and unsupervised feature extraction such as CP). We trained the model specifically to identify MOA, but the feature-extraction method can apply directly to any other discrete biological label of interest.

Certain considerations focus the scope of this study. First, MP performed poorly on certain MOAs (Supplemental Figure 5). This could be due to many reasons such as non-optimal experimental setups (e.g. with platemaps, drug concentration, cell type) or ineffective model architecture and training. Or, since Cell Painting only captured five channels, perhaps certain MOAs do not affect any of the biological markers used in the Cell Painting assay, and hence were phenotypically indistinguishable from control. Second, all the analyses were performed on compounds with just one known MOA. Understanding drugs that are associated with multiple MOAs is an important task, but our study did not address this question. Third, our study spanned just two concentrations: 5μM and 10μM. Generalizability to other concentrations, particularly more clinically relevant lower concentrations, has yet to be explored. Fourth, when we evaluated the method on completely held-out compounds (Figure 5), a minority of compounds had their MOAs correctly identified (at most 13.6% for JUMP1 and 10.7% for LINCS depending on the prediction method). Perhaps this is a limitation of Cell Painting’s ability to have distinguishable phenotypes for certain MOAs. Or perhaps compounds even within the same MOA class exerted unique phenotypic changes. For certain compounds, these changes may have been too different than what the model was exposed to during training. Performance on held-out compounds was still higher than both CP and DP (and multiple folds above a random classification: at most 23.9x for JUMP1 and 45.4x for LINCS). But perhaps performance can be enhanced by providing more compound diversity to the training set. We found that reducing the stringency from an exact match to a top-k match allowed for more compounds to be accurately predicted (25.4% of compounds for JUMP1 and 13.0% for LINCS for top-five MOA predictions). In practice, adopters of the method can probe the model’s top predictions during a compound’s experimental validation to increase the likelihood of discovering its true MOA. Lastly, practical considerations may also inform the application of the method. EfficientNets require high memory consumption and graphic processing unit (GPU) hardware for tractable training. However, if adopters of the method simply apply MP without any training on their datasets, then memory and compute constraints are much less limiting. If a user trains on custom datasets, we found the model did not need all technical replicates for near-maximal performance, especially with the LINCS dataset, indicating that identifying MOAs is possible in smaller datasets (Supplemental Figure 6).

Here we provide a tool for creating quantitative representations relevant to the task of MOA identification. With an MOA-specific embedder, we can query a drug’s phenotypic effect on cells and determine its MOA by similarity. We can then follow up with these predictions via traditional target-based screens. This strategy of broad profiling followed with target-based experiments can potentially be a powerful and cost-effective means of searching for new therapeutics. We hope that the method will be readily applicable to archival Cell Painting datasets and of broad use to future phenotypic screening endeavors. To facilitate open sharing, the model and source code are freely available at https://github.com/pfizer-rd/moa-profiler.

## Methods

### Dataset Preparation

The JUMP1 dataset is described in Chandrasekaran et al^30^. Download instructions can be found here: https://github.com/jump-cellpainting/2021_Chandrasekaran_submitted. Briefly, they conducted the Cell Painting protocol with three perturbation types: compound, CRISPR, and ORF. We kept only the compound data that had no more than one known MOA according to the CLUE Connectivity Map^35^, which can be found at https://clue.io/. Experimentalists used both U2OS and A549 cells, which were subjected to both 24 and 48 hours of compound treatment before being imaged. The resulting dataset after filtering single-MOA compounds consisted of 81,310 images (66,270 excluding DMSO) and 7,958 wells (6,486 excluding DMSO) coming from 23 384-well plates.

The LINCS dataset^32,39^ can be found at doi: 10.5281/zenodo.5008187. Briefly, the authors conducted a Cell Painting protocol with compound perturbation. Like the JUMP1 dataset, we kept only compound data that had no more than one known MOA according to the CLUE Connectivity Map. We also sub-selected compounds at 10μM concentration. Authors used only A549 cells, which were subjected to 48 hours of compound treatment before being imaged. The resulting dataset after filtering for single-MOA compounds at 10μM concentration consisted of 87,729 images (56,283 excluding DMSO) and 9,749 wells (6,255 excluding DMSO) coming from 136 384-well plates.

### Data Preprocessing and Model Training

We randomly split the dataset into a 60% training, 10%, validation, and 30% test split, such that each well and all its image fields were assigned to only one of the three sets. Each MOA was present in each of the three sets. We then class-balanced the training set so that each MOA had equal representation. We also included the negative DMSO as a class to learn but excluded it from all performance metrics because of its overrepresentation in the dataset.

For both datasets, images were captured in five channels at a resolution of 1080 × 1080px. The five channels came from five different biomarkers: Hoechst 33342 for DNA, Alexa 488 for the endoplasmic reticulum, Alexa 488 long for RNA, Alexa 568 for the actin cytoskeleton, golgi, and plasma membrane, and Alexa 647 for the mitochondria. We first scaled each image to the range zero to one, and then unit normalized them to a mean of zero and standard deviation of one. We permuted each channel’s brightness and contrast independently by a random factor in the range of 0 to 0.30 (just for the LINCS dataset). As a final training augmentation step, we performed random 90-degree rotations on each image, along with random horizontal flips. For a given field of view, we stacked each of the five augmented image channels into a single tensor.

We then fed this tensor (5 × 1080 × 1080) into a modified Efficientet-B0 architecture with the task of classifying the image’s compound’s MOA. The only modification to EfficientNet was adjusting the input layer to receive five channels instead of three. We trained for 100 epochs and selected the model that had the highest accuracy on the validation set. We used a learning rate of 0.1, a weight decay of 0.0001, a dropout rate of 0.2, a learning momentum of 0.9, a learning rate scheduler with a gamma decay of 0.1 at epoch 50 and 75, and batch size of 56 for training. We used four NVIDIA A-100 GPUs.

For training on smaller subsets of data (Supplemental Figure 6), all hyperparameter choices were kept the same as the training scheme on the full training set. The compound-replicate wells excluded from training were chosen randomly for each training run. The maximal allotted compound-replicates values went up to the maximal number of compound replicates in the full training set.

### CellProfiler and DeepProfiler Extraction

All CP embeddings were provided at the well-level (in this case, average embeddings of all single cells within a well). We downloaded CP embeddings for the JUMP1 dataset from https://github.com/jump-cellpainting/2021_Chandrasekaran_submitted. The CP pipeline used for extraction can be found at https://github.com/jump-cellpainting/2021_Chandrasekaran_submitted/tree/main/pipelines/2020_11_04_CPJUMP1. We downloaded CP embeddings for LINCS from: https://github.com/broadinstitute/neural-profiling/tree/main/baseline/01_data. https://github.com/broadinstitute/neural-profiling/tree/main/baseline The CP pipeline used for extraction can be found at https://github.com/broadinstitute/lincs-cell-painting/blob/master/profiles/README.md.

The repository for DP can be found at https://github.com/cytomining/DeepProfiler. DP leverages an EfficientNet trained on the ImageNet dataset for profiling. We derived DP embeddings using the “profile” function of DP with the default configurations and pre-trained weights automatically downloaded by DP. We aggregated single-cell DP embeddings into well-level embeddings by taking the median over all single-cell embeddings within a well.

When comparing MP and DP when both were allowed to train on the LINCS dataset (Supplemental Figure 4), we extracted all available DP embeddings from https://github.com/broadinstitute/neural-profiling/tree/main/training/runs/1102 and used them in the comparative analysis. We chose these embeddings because the author reported the highest validation accuracy on this set. Outlier control was not applied to any of the CP, DP, or MP embeddings.

### Classification Analysis

For all classifier evaluation metrics (Figure 3A, Figure 6A), we binarized the classes using sklearn’s label_binarize function, and plotted using sklearn’s precision_recall_curve function. Random baseline was set to the positive prevalence of the binarized labels. We did not include the negative control DMSO condition for analysis of raw-image classification due to its large class over-representation.

### Embedding Extraction and Analysis

Once the MP models were fully trained, we applied them to the held-out test set of wells never exposed to the model. To extract MP embeddings, we fed each image within a well through the EfficientNet, extracted the last convolutional layer, performed an average pool operation, and flattened the result. This image-field level embedding had a size of 1,280. We then took the median of all image-field level embeddings that belonged to the same well and assigned this as the well-level embedding.

Once we had well-level embeddings, we performed four analyses to assess how well the embeddings captured MOA-specific features. We excluded negative control DMSO wells from all embedding analyses.

First, we calculated the enrichment factor, which was the odds ratio in a one-sided Fisher’s exact test. This test assessed whether high PCC similarity for an embedding pair is independent of the embeddings sharing the same MOA. We used a range of PCC percentiles (from the 98^th^ to the 99.8^th^ percentile) as the thresholds for determining which embedding pairs were considered strongly correlated (versus weakly correlated). The odds ratio was calculated as (a/b) / (c/d) for the following 2×2 frequency table:

|  | MOA replica | Non-MOA replica |
| --- | --- | --- |
| Strongly correlated | a | b |
| Weakly correlated | c | d |

Second, we calculated the average pairwise PCCs for two groupings of well-level of embeddings: those that had the same MOA, 2) those that had different MOA. Within each grouping, we calculated the PCC for each pair of wells and averaged the result. Delta values were the differences between the two groups’ averages.

Third, we calculated k-NN metrics by finding the k closest neighbors for each well based on PCC similarity between the well embeddings, and taking the majority MOA of those neighbors as the predicted MOA. We evaluated all possible values of k from one to the total number of embeddings minus one. For each k, we used sklearn’s f1_score, precision_score, and recall_score functions. Averages were weighted by support (the number of true instances for each label).

Fourth, we derived a MOA-level embedding (MLE) by grouping all the wells of that MOA and taking the median. We treated each well in the test set as a query well. For each query well, we calculated the class embeddings with the query well excluded, and then assigned a prediction for the query well’s MOA based on which MLE was the most similar (by PCC) to the query well’s embedding. We calculate weighted-average F1, precision, and recall scores for these predictions with a one-vs-rest scheme using the sklearn’s f1_score, precision_score and average_recall_score functions.

As a last metric of embedding integrity and MOA-specificity, we calculated pairwise PCC averages of three groups: 1) embeddings with the same compound (and hence same MOA), 2) embeddings with different compound but same MOA, and 3) embeddings with different compound and different MOA. All pairs were unique and an embedding was never paired with itself. We determined statistical significance of difference of means with a two-sided z-test (see “Statistical Tests”).

### Analysis for MOAs Represented by Multiple Compounds

For Supplemental Figures 1 and 2, we analyzed only MOAs that were represented by multiple different compounds (at least two unique compounds for each MOA). Single-compound MOAs and all of their corresponding images were filtered out of the test set. For this smaller subset, we kept the same analysis as the one with all available MOAs (Figures 3 and 6).

### Analysis for Held-out Compounds

For the held-out compound analysis (Figure 5), we split the dataset by compound instead of by well. We held out one randomly selected compound for each MOA class that was represented by at least two unique compounds. All other compounds were used for training and validation, with 70% of the wells used for training, and 30% for validation. We trained a new model based on this dataset split using the same hyperparameter choices as the analysis for the well-split scheme. We evaluated two different ways to make compound MOA predictions: using the model’s classification outputs and the model’s latent representations. We analyzed both to account for two circumstances of probing an unknown compound: 1) if an adopter of the method would like a discrete MOA label (within the set of MOAs in this study), or 2) if an adopter would like a more general latent representation for a compound.

1. We used the model’s classification outputs (i.e. the resulting classification vectors of the EfficientNet’s final linear layer). We derived a single MOA prediction for each image field. For each well, we assigned the well’s MOA prediction as the majority MOA prediction over its image fields. Finally, for each compound we predicted its MOA as the majority MOA prediction over its wells. We varied the stringency of this prediction and computed different accuracies for different values of k such that the held-out compound is correctly predicted if the correct label is among the top-k MOA well predictions. MOAs in the set of k=n included all MOAs with the highest n counts. K spanned from k=1 (strictest exact match between top prediction and true label) to k=5.
2. We used the model’s latent space representation embeddings. For the training set, each MOA was assigned a MLE by aggregating (via median) over all well-level embeddings belonging to the MOA.

a. Individual vote: For each well-level embedding in the held-out test set, we assigned its MOA prediction as the MOA of the most similar MLE via PCC. Finally, we assigned a compound’s MOA prediction as the majority MOA over these well-level predictions.
b. Aggregated vote: For each held-out compound, we derived an aggregated compound-level embedding (CLE) by taking the median over all well-level embeddings of the compound. Finally, we assigned the compound’s MOA prediction as the MOA of the MLE most similar to CLE by PCC.

### Statistical Tests

For all statistical test of significance, we performed a two-sided z-test for difference of means. We used a null hypothesis stating that the means were equal, and an alternative hypothesis stating that the means were different. Significance was set at *P* < 0.05. Each sample corresponded to a well-level embedding.

