## Supplement for "Deep Representation Learning Determines Drug Mechanism of Action from Cell Painting Images"

1. *Machine Learning and Computational Sciences, Pfizer Worldwide Research Development and Medical, 610 Main Street, Cambridge, Massachusetts 02139, United States*
2. *Internal Medicine Research Unit, Pfizer Worldwide Research Development and Medical, 610 Main Street, Cambridge, Massachusetts 02139, United States*
3. *Discovery Sciences, Pfizer Global Research and Development, Groton Laboratories, 280 Shennecossett Rd, Groton, CT 06340*

### Supplemental Figure 1

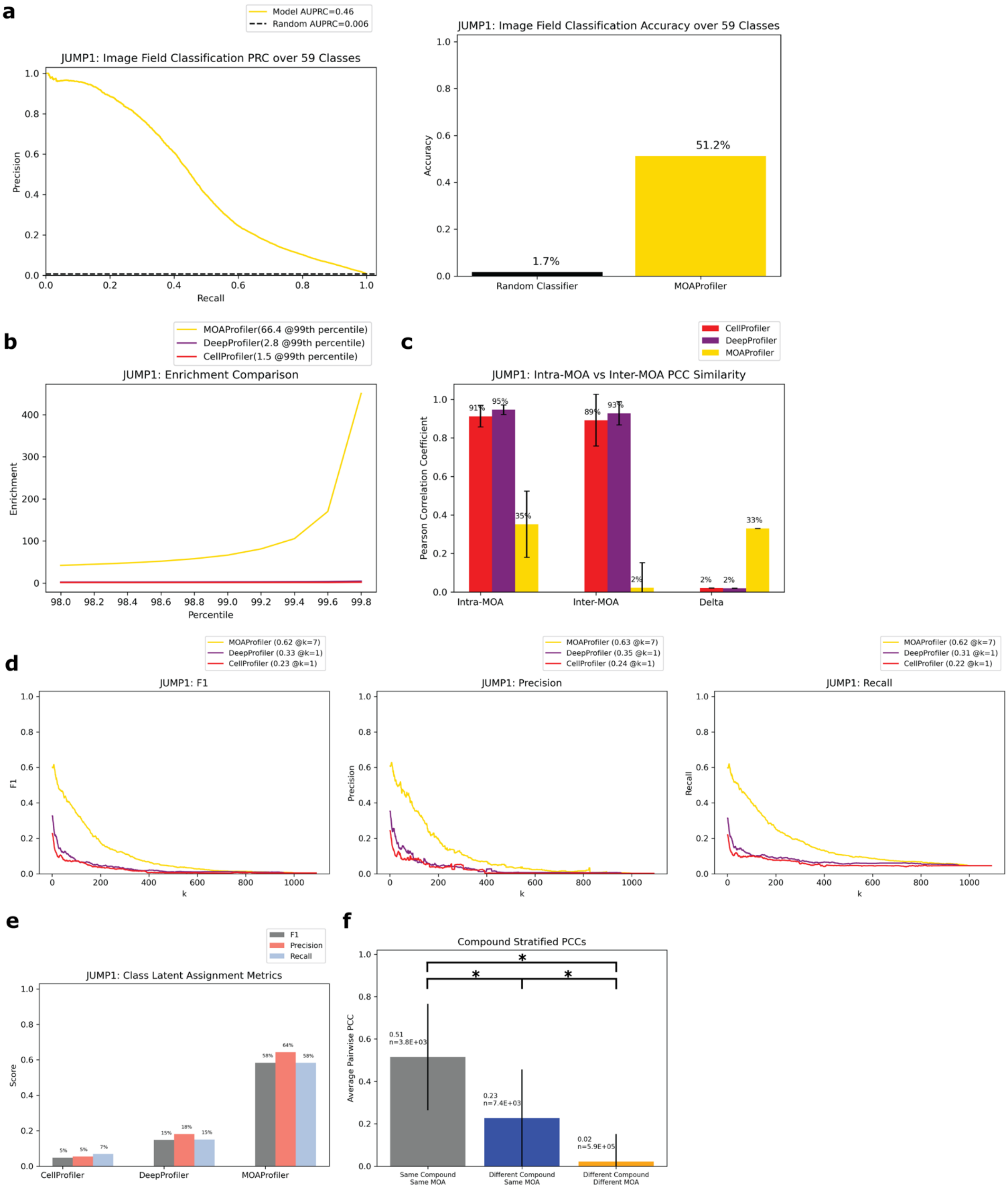

**Supplemental Figure 1: Performance on a JUMP1 subset that only included MOAs each represented by more than one unique compound.** This resulted in 59 unique MOAs. Otherwise, analysis was equivalent to Figure 3.

a) PRC and accuracy of image classification. Random baseline for PRC is 1 / 59.
b) Enrichment comparison at different percentiles.
c) Intra-MOA vs inter-MOA average pairwise PCC similarity.
d) K-NN embedding metrics.
e) Class latent assignment metrics.
f) Average pairwise PCCs of three different groups of pairwise embeddings stratified by perturbation.

### Supplemental Figure 2

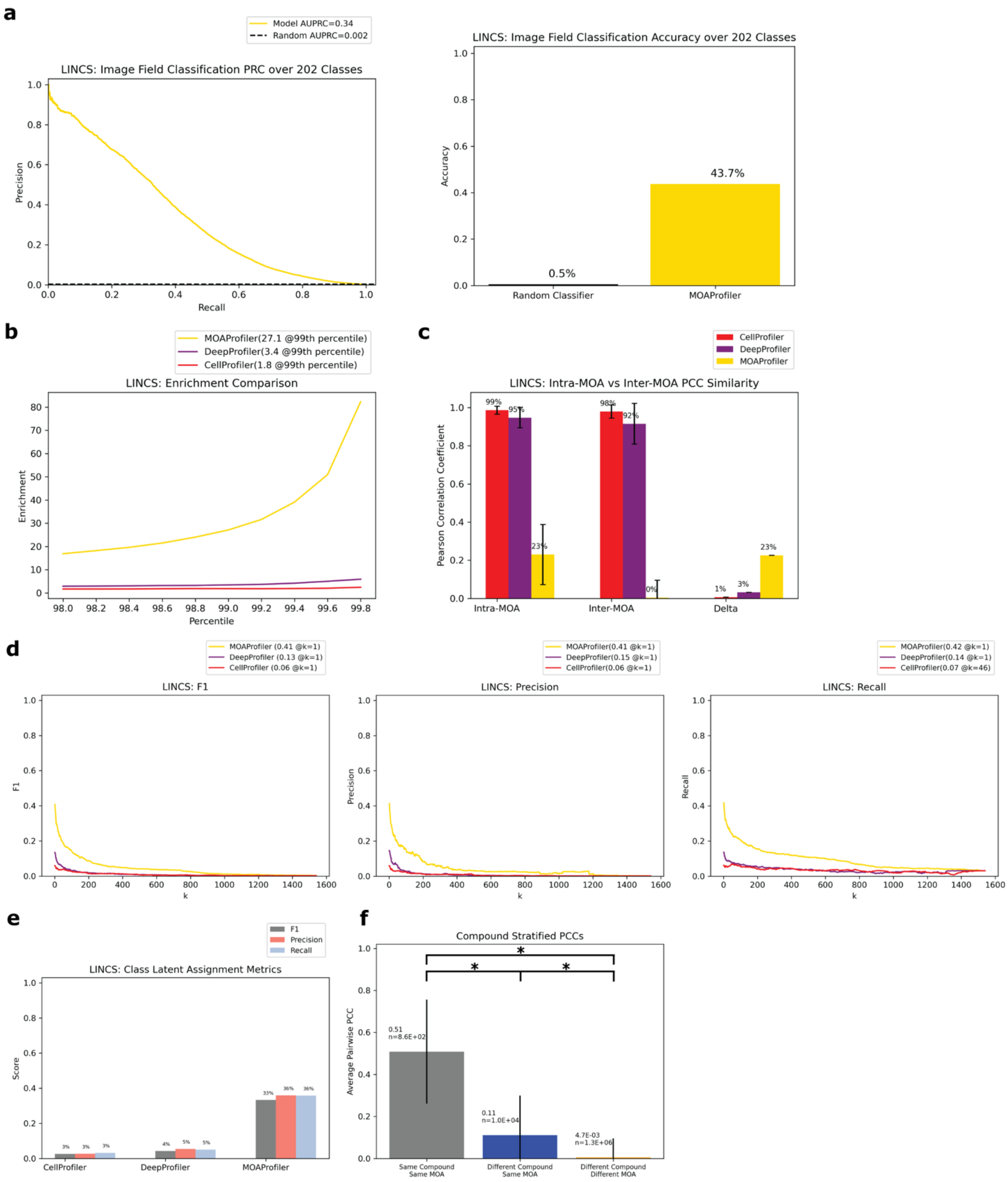

**Supplemental Figure 2: Performance on a LINCS subset that only included MOAs each represented by more than one unique compound.** This resulted in 202 unique MOAs. Otherwise, analysis was equivalent to Figure 6.

a) PRC and accuracy of image classification. Random baseline for PRC is 1 / 202.
b) Enrichment comparison at different percentiles.
c) Intra-MOA vs inter-MOA average pairwise PCC similarity.
d) K-NN embedding metrics.
e) Class latent assignment metrics.
f) Average pairwise PCCs of three different groups of pairwise embeddings stratified by perturbation.

Supplemental Figure 3

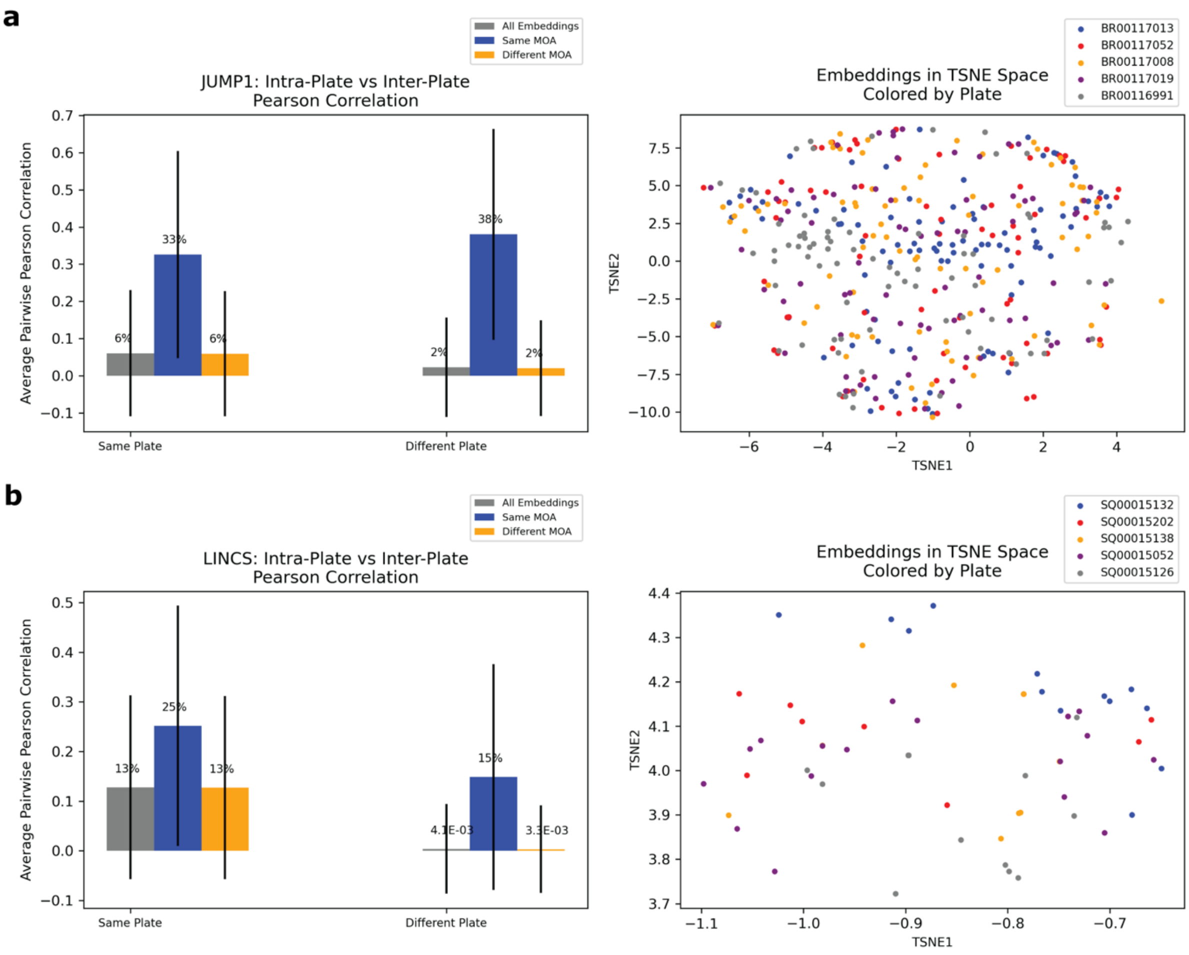


**Supplemental Figure 3: Quantifying batch effects on model embeddings.** a) JUMP1 b) LINCS. Left: average pairwise PCCs over the test set for three groups: 1) embeddings from different plates (grey), 2) embeddings from different plates with the same MOA (blue), and 3) embeddings from different plates with different MOAs (orange). Error bars span one standard deviation in each direction. Right: TSNE visualization of all test-set embeddings from five randomly chosen plates. Embeddings colored by plate.

### Supplemental Figure 4

**a**

| MOA Profiler | Deep Profiler |
|---|---|
| Does not require single cell locations | Requires single cell locations |
| Training classification at field level | Training classification at single-cell level |
| gamma decay learning rate scheduler | cosine learning rate scheduler |
| Strong supervision: classify MOAs, extract embeddings for MOA determination | Weak supervision: classify compounds, extract embeddings for MOA determination |
| No label smoothing | Label smoothing |

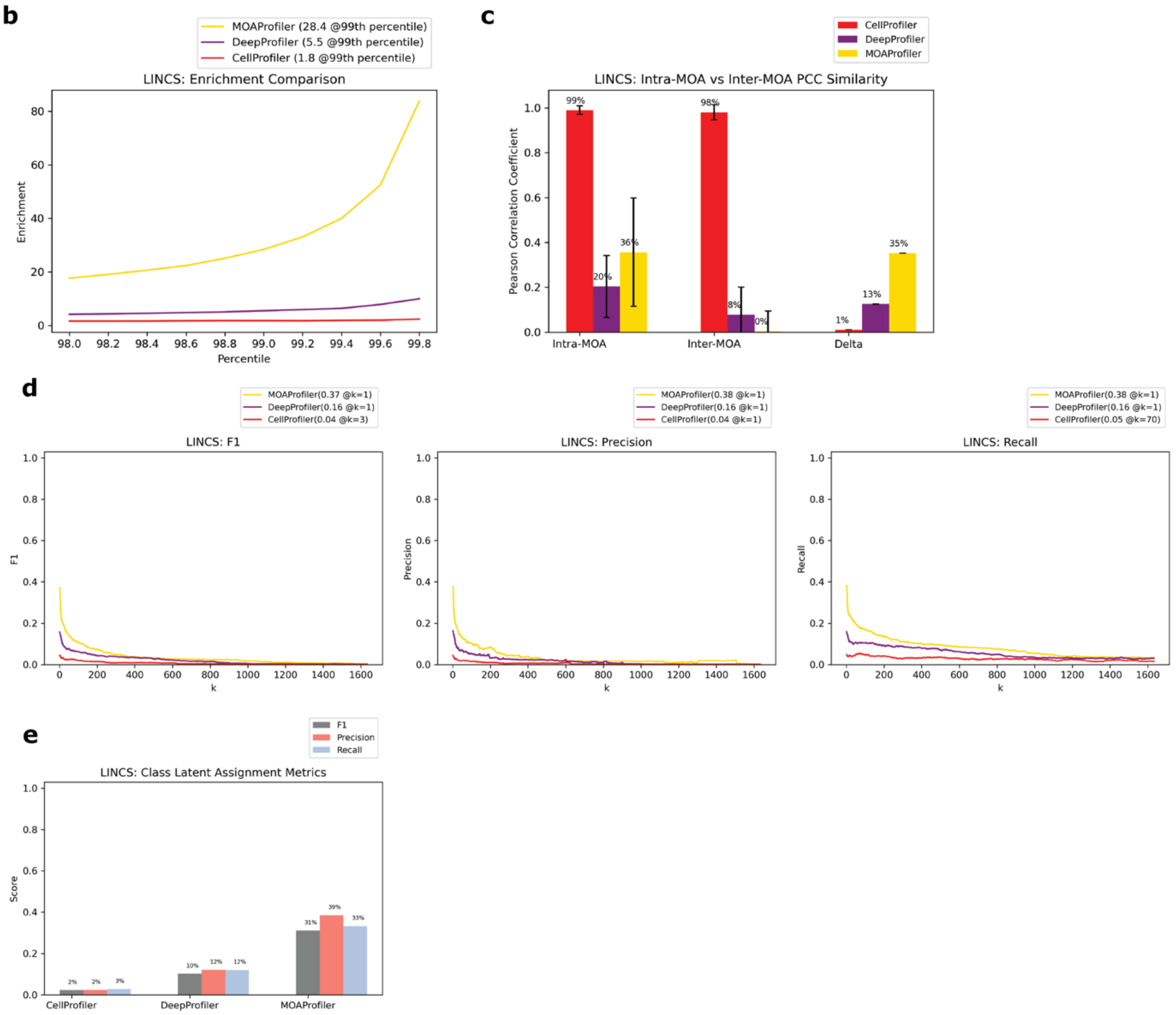


**Supplemental Figure 4: Comparing MP to DP both trained on the LINCS dataset.** Otherwise, embedding analysis was the same as Figure 6.

a) Key differences between MP and DP methods for MOA determination.
b) Enrichment comparison at different percentiles.
c) Intra-MOA vs Inter-MOA average PCC similarity.

d) k-NN embedding metrics.
e) Class latent assignment metrics.

Supplemental Figure 5

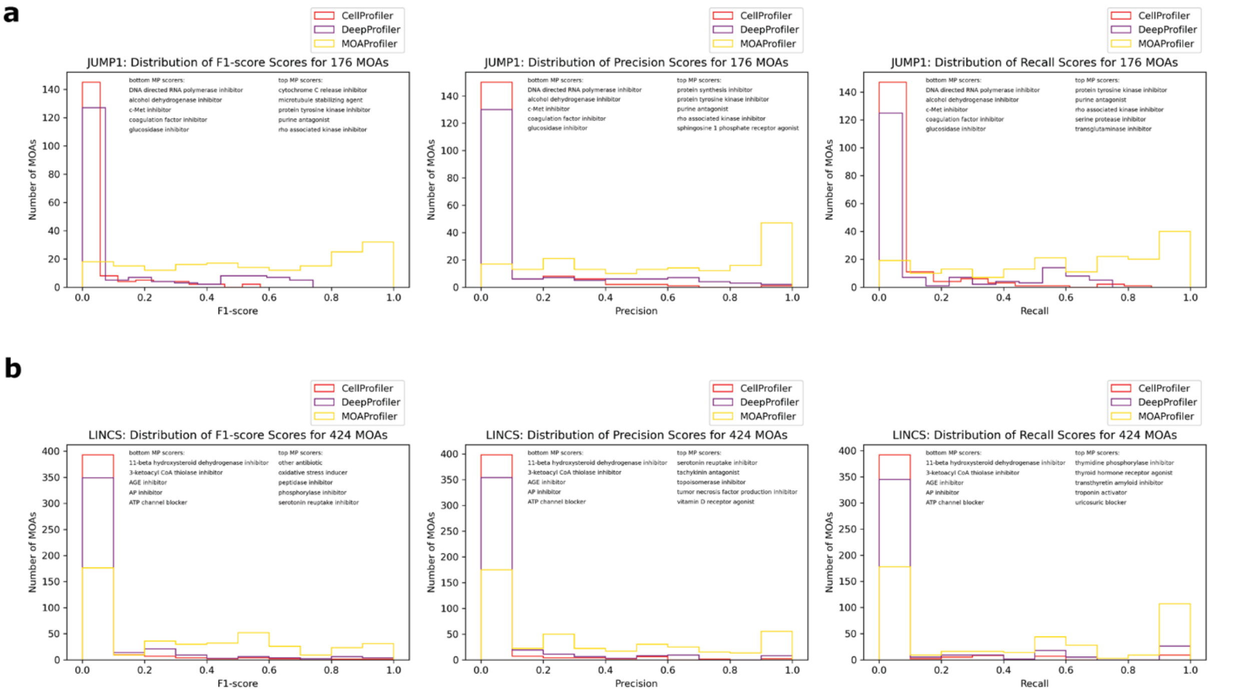


**Supplemental Figure 5: Distribution of class latent assignment scores over the test set for each MOA.** a) JUMP1 b) LINCS. X-axis: score, y-axis: count of MOAs achieving the class latent assignment score. Listed: worst and best performing MOAs by score after being embedded by MP.

### Supplemental Figure 6

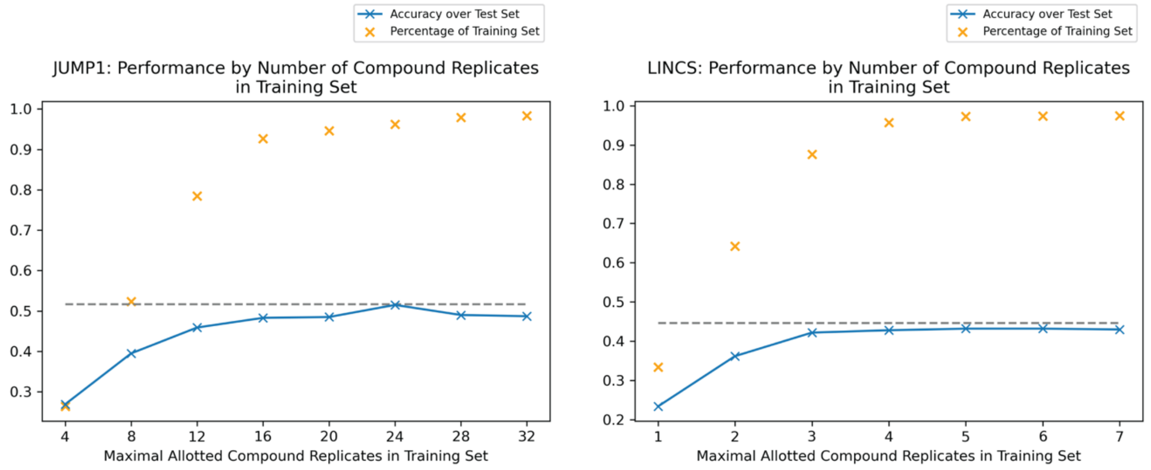


**Supplemental Figure 6: Performance with smaller training sets.** JUMP (left), LINCS (right). Keeping the validation and test sets the same, we systematically limited the number of compound-replicate wells present in the training set and trained new models from these smaller sets (Methods). X-axis: maximal number of compound-replicate wells allowed in the training set, blue: accuracy across held-out test set, orange: percentage of images compared to the full training set. Dashed horizontal line indicates the performance over the test set when keeping the full training set.
